# Predator cues and environmental enrichment during development drive intraspecific variation in the behaviour and life history of juvenile lobsters

**DOI:** 10.1101/2024.02.10.578266

**Authors:** Giovanni Polverino, Lorenzo Latini, Giuseppe Nascetti, Giacomo Grignani, Eleonora Bello, Claudia Gili, Claudio Carere, Daniele Canestrelli

**Affiliations:** Department of Ecological and Biological Sciences, University of Tuscia, Viterbo, Italy; School of Biological Sciences, Monash University, Melbourne, Australia; School of Biological Sciences, University of Western Australia, Perth, Australia; Stazione Zoologica Anton Dohrn, Naples, Italy

**Author notes:** These authors contributed equally: Claudio Carere and Daniele Canestrelli. Joint first authors (Giovanni Polverino, Lorenzo Latini). Corresponding author, Postal address: Viterbo, 01100 VT, Italy.

**Keywords:** Animal personality, Behavioural plasticity, Conservation aquaculture, Crustaceans, Developmental plasticity, Phenotypic plasticity

## Abstract

Intraspecific variation in ecologically relevant traits is critical for animal populations to survive in our rapidly changing world. This is especially true for species that suffer from intense harvesting regimes, whereby populations density is often low. Standard hatchery procedures can assist some management and conservation programs by producing large numbers of juveniles to be released into the wild. Yet we know surprisingly little on the impact that such standard, minimalistic settings have on the development of intraspecific variation in important phenotypes of the individuals, including among-individual variation in behavioural (individuality) and life-history traits, and in the plasticity of those traits in response to varying environmental conditions. Here, we fill this gap by testing whether early-life exposure to different environmental conditions alters the development of individuality and plasticity in ecologically relevant behaviours and life-history traits of the European lobster (*Homarus gammarus*)—one the most harvested species in the Mediterranean, which has been subjected to conservation programs for decades. By accessing one of the largest lobsters hatcheries in Italy, we used the progeny of wild-caught females and manipulated—in a full factorial design—the environmental complexity of the individual enclosures (i.e. presence/absence of substrate and/or shelter) and the level of exposure to cues from their natural predators. We repeatedly quantified behaviours (i.e. activity, refuge use, and aggressiveness) and life-history traits (i.e. carapace length and intermoult period) of the individuals throughout their early development, capturing both mean and individual-level variation across treatments. Our results offer solid evidence that effects of standard hatchery settings extend far beyond mean changes in the behaviour and life history of the animals, compromising the development of individual plasticity in those traits that are essentials for populations to survive in the wild—likely reducing the effectiveness of conservation programs.

## INTRODUCTION

In conservation aquaculture, aquatic organisms are typically produced in captivity in large numbers for their reintroduction into the wild, with the aim of restoring depleted populations and enhance wild stocks (Froehlich et al., 2017). However, conventional hatchery-rearing procedures impose substantially different selection pressures on the individuals relative to their natural environment, potentially altering the phenotypic traits that are critical for these organisms to survive in the wild (McDougall et al., 2006; McPhee & McPhee, 2012).

Mounting evidence suggests that changes in the phenotypic composition of a population and in the ability of captive-reared individuals to adjust to changing conditions of their environment can impact the ultimate outcome of wildlife management and conservation programs (McDougall et al., 2006; Mason, 2010; Crates et al., 2023). For instance, phenotypic differences among the individuals are known to be essential for animal populations to survive in our increasingly changing world (Sih et al., 2004; Dall et al., 2012; Polverino et al., 2021). A large portion of such phenotypic diversity within animal groups can be attributed to the effect of different environmental conditions experienced by the individuals (Stamps, 2015), and is quantified as individual-level variation in mean traits and in their plastic adjustments over time and/or across contexts (Dingemanse & Dochtermann, 2013; Snell-Rood, 2013). Nevertheless, while growing evidence suggests that phenotypic variation among individuals can drive evolutionary changes and sustain functional diversity across populations, communities, and ecosystems (Miller & Rudolf, 2011; Des Roches et al., 2018; Raffard et al., 2018), a controversy remains regarding the proximate and ultimate mechanisms of such variation (Polverino et al., 2016).

Early-life conditions experienced by an individual are known to play a key role in shaping the development of its phenotype in adulthood (Monaghan 2008; Frankenhuis & Panchanathan, 2011), and so its capacity to cope with environmental challenges. Yet environmental effects during ontogeny are usually overlooked as the proximate mechanisms behind the emergence of individual differences compared to environmental effects operating across generations (i.e., genetic adaptation; Urszán et al., 2015 and 2018). It is therefore important to understand whether and how environmental conditions can contribute, at least in the short term, to the emergence of phenotypic variation at the individual level. For example, other than improving animal welfare conditions (Zhang et al., 2021), introducing ecological-relevant cues into the hatchery-rearing environment could also benefit conservation programs, by promoting diverse coping strategies and larger investments into phenotypic plasticity (Shepherdson, 1994; Watters & Meehan, 2007; Reading et al., 2013; Daly et al., 2021). Thus, shedding light on these aspects is critical for managing risks, ensuring the effectiveness of rearing programs, and guarantee high standard of welfare conditions (Webster, 1995; McDougall et al., 2006).

Here, we wanted to test whether and how the early-life exposure to different environmental conditions altered the development of ecologically relevant behaviours and life-history traits of hatchery-reared juvenile European lobsters (*Homarus gammarus*): an ecologically and economically important decapod species, historically targeted for conservation programs. Specifically, 256 lobsters were exposed to different degrees of environmental complexity during their early benthic stages, which included the presence/absence of structural enrichments into their individual rearing enclosures (shelter and substrate) and cues from natural predators. Juveniles were repeatedly assayed during their early development for ecologically relevant behavioural (activity, refuge use, aggression; Réale, 2007) and life-history traits (carapace length and intermoult period; Chang et al., 2012). This approach allowed us to test for treatment effects on both mean and individual-level variation in behaviour and life history (Dingemanse et al., 2010). In accordance with previous studies (Aspaas et al., 2016; Arechavala-Lopez et al., 2020), we hypothesized that standard, minimalistic aquaculture settings would compromise the behavioural repertoire of the animals and alter their growth. Furthermore, we expected that effects of standard hatchery conditions on lobsters’ development would extend beyond mean changes in their behaviour and life history, and impair also the capacity of the individuals to adjust their phenotypes to rapid changes of their surrounding environment (e. g. presence of predators). Our analysis sheds light onto the role that early-life environmental stimuli have in shaping mean and individual-level variation in animal phenotypes, offering important insights for science-based improvements of aquaculture practices and conservation programs.

## METHODS

### Study species and experimental animals

We used first-generation progeny of wild-caught European lobsters (*Homarus gammarus*), which were hatched under controlled conditions in the Ichthyogenic Experimental Marine Centre (CISMAR) at the University of Tuscia (Tarquinia, Italy; 42.201851, 11.721749). Four ovigerous females were caught by local fishermen from two distinct locations along the Tyrrhenian coast (Porto Ercole: 42.393183, 11.211192; Montalto Marina: 42.327235, 11.572900) and transported to the CISMAR facilities. The females were housed separately into 1500L tanks connected to a recirculating aquaculture system, which supplied seawater to the apparatus during the five months until the eggs hatched. The seawater flowed through a suspended solid removal unit (63µm mesh), a bio-filter reactor, a foam fractionator, and a UV sterilizer before entering the holding tanks. Water temperature was set at 17°C, while salinity, dissolved oxygen, pH, and nitrogen (NO_2_^-^ and NO_3_^-^) were monitored daily and kept within the optimal range for the species (Hinchcliffe et al., 2021).

Eggs were monitored daily and the newly hatched (planktonic) larvae were collected every morning, counted, and housed separately for each female into 200L upwelling, aerated vessels at low stocking density (∼20 larvae L^-1^; Hughes et al., 1974; Bell et al., 2005; Hinchcliffe et al., 2021). The vessels were connected to the same recirculating system described above, so that water conditions experienced by the newborns reflected conditions during their incubation. Larvae were fed *ad libitum* twice a day with a mix of frozen *Artemia* sp., *Mysis* sp., and krill (*Euphasiidae* sp.).

### Experimental procedure

Approximately 14 days after birth the larvae reached the benthic stage. Then, 256 juveniles (64 per female) were randomly selected for the experiment and distributed across the experimental treatments, in which the animals were maintained for 24 consecutive weeks (i.e. 168 days). Specifically, four identical floating grids were set up for the experiment, each one placed inside a 1500L tank. Each grid consisted of 64 individual squared-shaped enclosures (8 cm per side and 3 cm deep each), which were made of solid plastic material with a perforated bottom surface (1 mm^2^) to ensure the adequate water exchange between the individual enclosures and the housing tank. For each grid, individual tanks were randomly split across four rearing treatments that differed from each other for the presence/absence of a substrate (calcium carbonate gravel) and/or a shelter (PVC pipe): no substrate and no shelter (control), substrate, shelter, and substrate and shelter (Fig. 1a).

**Figure 1.**
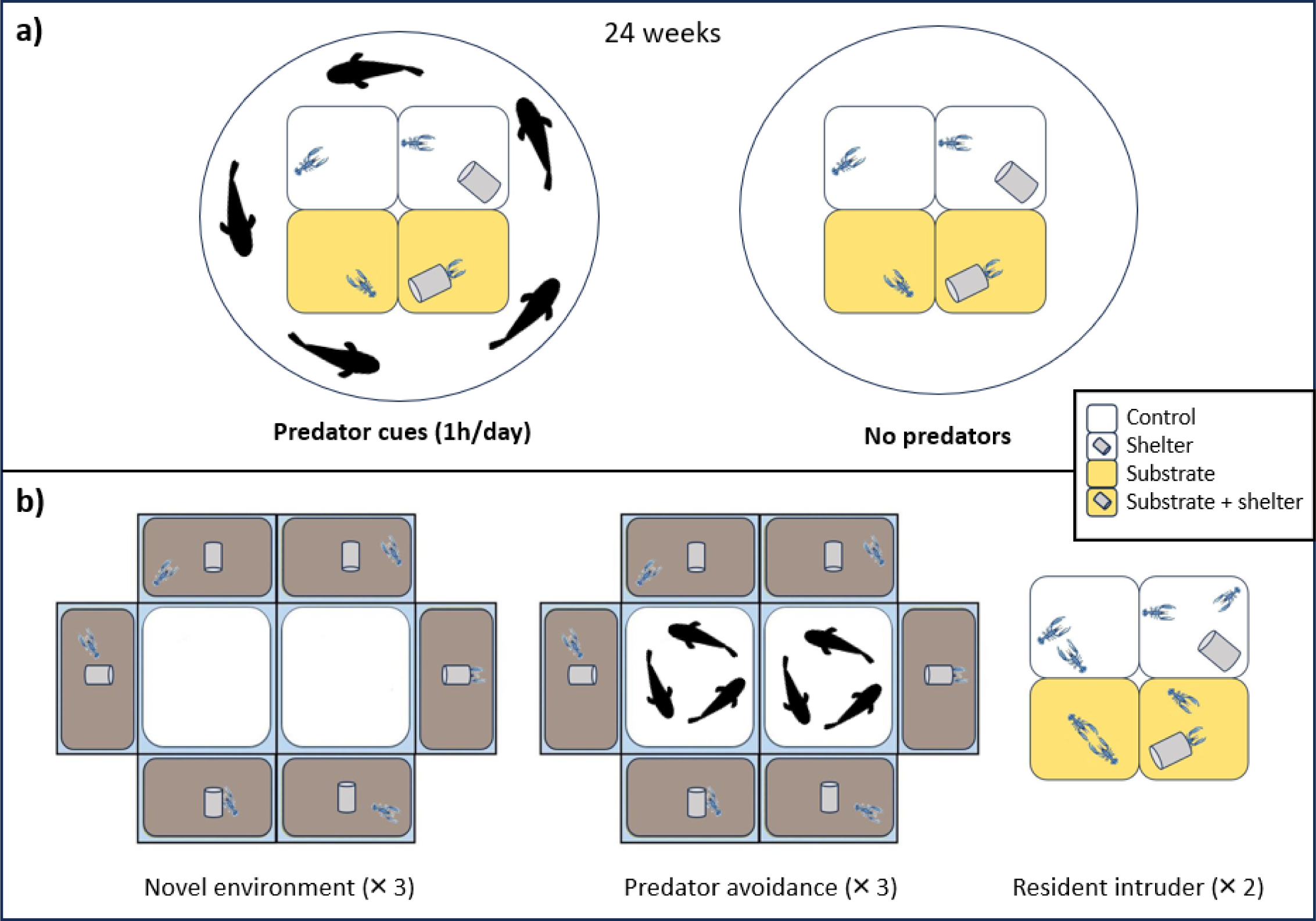
Representation of the experimental procedure. **a)** Ontogenetic treatments. Juveniles raised in individual compartments were exposed to different rearing treatments that differed for the presence/absence of environmental stimuli: shelter, substrate, and predator cues (1h per day). Juveniles were subjected to these conditions from the beginning of their benthic life for approximately 24 consecutive weeks. **b)** Acute treatments and behavioural tests: novel environment and predator avoidance. The setup consisted of two squared, central areas to house the predator during the predator avoidance test, and six visually isolated rectangular compartments (lobster’s tanks) equipped with substrate and a shelter. The resident-intruder test was performed at the seventh benthic stage on a subsample of the animals.

Individuals from two of the floating grids were also exposed to predator cues, for one hour a day, throughout the 24 experimental weeks (predator cues treatment; Fig. 1a). On the contrary, animals housed in the other two grids were never exposed to predators (no predator treatment; Fig. 1a). Each tank containing a grid assigned to the predator treatment hosted five juvenile sea breams (*Sparus aurata,* 10 ± 1 cm in length)—a common predator of small crustaceans (Taieb et al., 2013)— for one hour per day. Individuals in the predator treatment could perceive the presence of their predator via chemical cues (Derby & Sorensen, 2008), but the opaque, perforated material of the individual enclosures prevented visual interactions between individuals and their predators. After the one hour of exposure, predators were removed and the grids with the experimental lobsters transferred into clean holding tanks. Control grids were also moved into different, clean holding tanks once a day, to balance the procedure across treatments.

Animals were fed daily with a mixture of frozen *Artemia* sp., *Mysis* sp. and frozen krill (*Euphasiidae* sp.) *ad-libitum*. The illumination reflected the light:dark cycle (8:30am-5:30pm) and was diffused through 36W aquarium lamps.

### Behavioural assays

The behavioural responses of the juveniles were scored, repeatedly over their ontogeny, across three different behavioural tests: novel environment (NE), predator avoidance (PA), and resident-intruder (RI, Fig. 1b). Behavioural tests were always performed one week after the animals moulted to standardize the procedure, as suggested by Tamm and Cobb (1978). To do so, the days between moults were monitored per each individual throughout the experimental campaign.

The NE and PA tests were performed on each animal consecutively in an experimental arena, in this order, to minimize manipulation, for a total of three replicates per individual (approximately at 3, 6 and 10 weeks post hatching). Instead, the RI test was repeatedly performed, twice on each individual, within each individual’s rearing enclosure.

Behavioural trials were recorded with a GoPro Full HD camera (48 frame per sec). Behavioural videos were scored, blind to the treatment, with the software Boris 5.1.3 (Friard & Gamba, 2016), to obtain latencies, duration, and frequencies of behavioural traits.

### Novel environment and predator avoidance test

The experimental arena consisted of two central, squared tanks (21.5 cm per side and 11 cm high; predator areas) surrounded by six rectangular test tanks (21.5 cm long, 11 cm wide, and 11 cm high), each one equipped with gravel substrate and a shelter (Fig. 1b). Before a novel environment test (NE) was initiated, two operators collected six experimental animals and placed them individually within the test tanks inside their shelters (Fig. 1b). After one minute, the shelters were simultaneously removed, and the juvenile lobster were free to explore the novel environment for a total of 5 min.

Soon after the test ended, three juvenile seabreams (10 ± 1 cm in length) were released into each of the two squared, central tanks, and the predator avoidance test (PA) was initiated. The behaviour of each individual lobster was recorded for further 5 min while in the presence of the predators (Fig. 1b). Notably, transparent and perforated walls divided the predator arenas from the lobsters’ test tank, so that lobsters were able to both visually and chemically perceive the presence of their predators, which is known to increase the perceived predation risk in prey animals than when they are exposed to simpler, unimodal cues (Ward & Mehner, 2010). At the end of the PA test, the juvenile lobsters were moved back into their housing containers and the test tanks were emptied and filled with clean water before the next trial commenced.

We chose activity (time spent moving, in sec) and refuge use (time spent inside the shelter, in sec) as the reference traits (Polverino et al., 2018 and 2021), since individual-level variation in these behaviours is a target of selection in non-sessile animals (Réale et al., 2007) and have ecological and evolutionary consequences (Wolf & Weissing, 2012).

### Resident-intruder test

The resident-intruder test (RI) was performed on 96 of the experimental individuals (randomly selected and balanced across treatments) once they reached the seventh benthic stage, that is, approximately 26 weeks since hatching (Fig. 1b). Individuals were assayed in their individual rearing containers, to test for their territoriality (Karnofsky & Price, 1989). To do so, 50 juvenile lobsters were used as the intruders, and were comparable in age and size to the focal individuals.

Before a RI trial started, a selected intruder was placed inside a perforated transparent cylinder located in the middle of the compartment. After 20 sec the transparent cylinder was gently lifted, and the intruder was free to physically interact with the resident lobster for up to two consecutive min (Fig. 1b). To minimise distress in the animals during the trials, a test was terminated soon after the first attack was observed. Trials were videorecorded with a GoPro Full HD camera (48 frame per sec) located 50 cm above the grid. Subsequently, the latency of the resident lobster to attack the intruder was recorded. Each intruder was changed after each trial, and a new intruder was randomly chosen so that focal animals never encountered the same intruder twice. Each focal individual was assessed twice, with tests four days apart.

### Morphometric measurements

Each juvenile was photographed with a Fujifilm xp140 camera and its carapace length (CL, mm) was measured with the dedicated software imageJ (Schneider et al., 2012), blind to the treatment, before each round of behavioural tests. Therefore, we collected three morphometric datapoints per individual. Additionally, we determined the period of time between two consecutive moults (i.e. intermoult period, IP, days) per each individual.

### Data analysis

A final sample size of 250 juvenile lobsters (control: n=63; substrate: n=61; shelter: n=62; substrate + shelter: n=64) out of 256 individuals completed all behavioural and morphometric assays—six juveniles were lost due to early mortality—for a total of 1.536 behavioural observations. Instead, 96 individuals performed the RI test, for a total of 192 behavioural observations. Overall, our sample size relied on > 130 h of video recordings.

Data analysis was performed with *RStudio* v. 4.2.1 (*R* Core Team, 2016), using the packages *lme4, lmerTest,* and *emmeans* (Bates, 2014; Kuznetsova et al., 2017; Lenth et al., 2023). We assumed Gaussian error distribution, which was confirmed for all response variables after visual inspection of model residuals. The significance level was set at α < 0.05.

We were interested to test whether and how the presence of environmental stimuli during development altered mean behaviours and life-history traits of the animals, as well as their group-level plasticity. To do so, we fitted generalized linear mixed-effects models with activity (i.e. time spent moving, in s), refuge use (i.e. time spent inside the shelter, in s), body size (i.e. carapace length, in mm), and intermoult period (i.e. time between two consecutive moults, in days) as the dependent variables, respectively. In the fixed-effect structure of the models for activity and refuge use we included rearing treatment (presence/absence of a shelter and/or substrate; four levels), predator treatment (predator cues and no predators), test (novel environment and predator avoidance), grid (four levels), mother ID (four levels), stage (three levels), test duration (in sec), and the interaction between rearing treatment, predator treatment, and test. In the aggressiveness model we included rearing treatment, predator treatment, their interaction, grid, and mother ID as the fixed effects. The models for carapace length (CL) and intermoult period (IP) included, instead, rearing treatment, predator treatment, stage, their (three way) interaction, grid, and mother ID as the fixed effects. We ran pairwise comparisons with the conservative Bonferroni method for significant predictors, while accounting for the variation explained by other predictors.

We included individual identities (random intercepts) in the random structure of all models to account for repeated measures. The aggressiveness model also included intruder identities as random intercepts. Since individuals might differ in the way that they adjusted their activity and refuge use across tests, developmental stages, and treatments or as function of their mother ID, we also tested for the presence of heterogeneous variance between tests, developmental stages, rearing treatments, predator treatments, and mother IDs (that is, random slopes/regression). We did this by running five separate models for activity and refuge use in which we included random intercepts and slopes one-by-one. Then, we used both likelihood ratio tests (LRTs) and Akaike information criteria (ΔAIC) to compare models with different random intercepts and slopes and chose models with the best likelihood ratio, and lower AIC. With the same approach, we also tested whether random intercepts explained a significant portion of the variation observed, that is, whether individual variation in activity and refuge use was highly structured (Dingemanse & Dochtermann, 2013).

## RESULTS

The rearing conditions to which individuals were exposed during their early development had a strong effect on the activity levels and aggressiveness of the animals, with marginal, albeit non-significant, effects also on the growth rate of the individuals (i.e. intermoult period; Table 1). In particular, the presence of both substrate and a shelter in the housing enclosure resulted in individuals being more active than their siblings raised under both control conditions and in the presence of substrate only (Fig. 2a). Instead, juveniles raised with both substrate and shelter, or with substrate only, were less aggressive toward intruders (i.e. longer latencies to attack) than conspecifics raised under control conditions (Fig. 2c). On the contrary, no significant effects of rearing conditions were observed for refuge use and carapace length (Fig. 2b).

**Figure 2.**
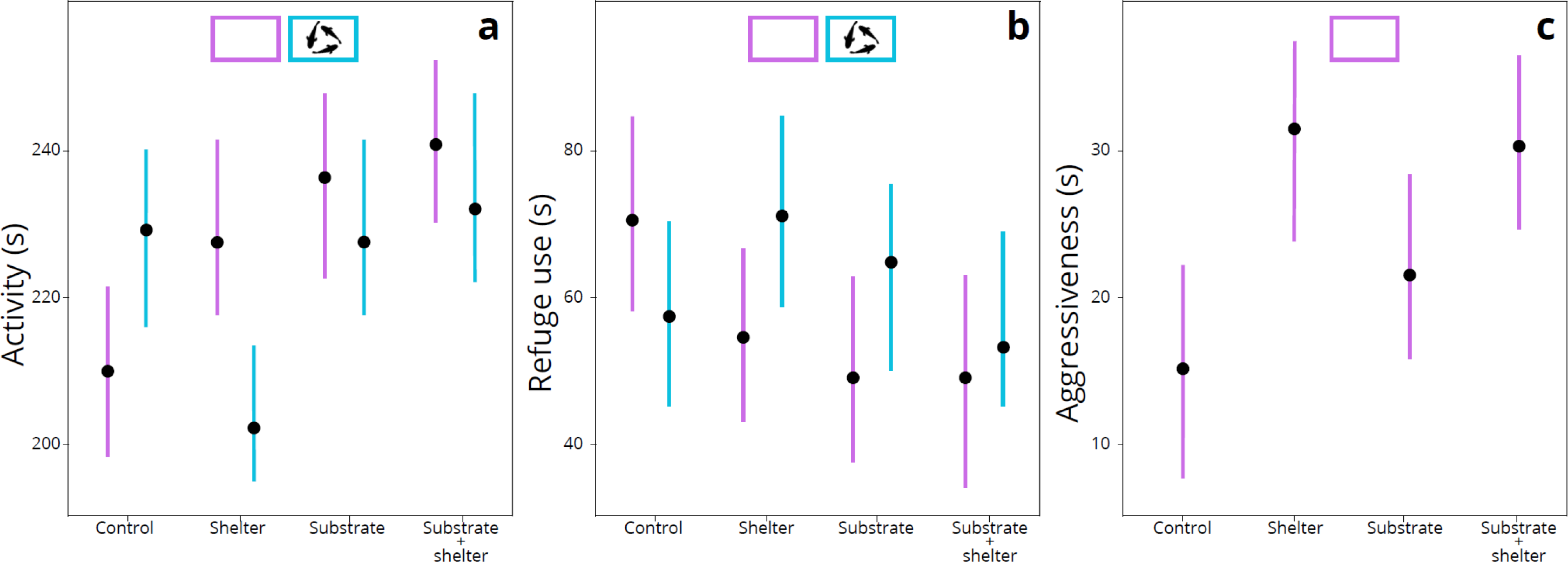
Effects of the different rearing conditions on the expressed behaviours. In (a) and (b) are respectively represented the treatment effects on activity and refuge use, and in (c) the effect on aggressiveness.

**Table 1.**
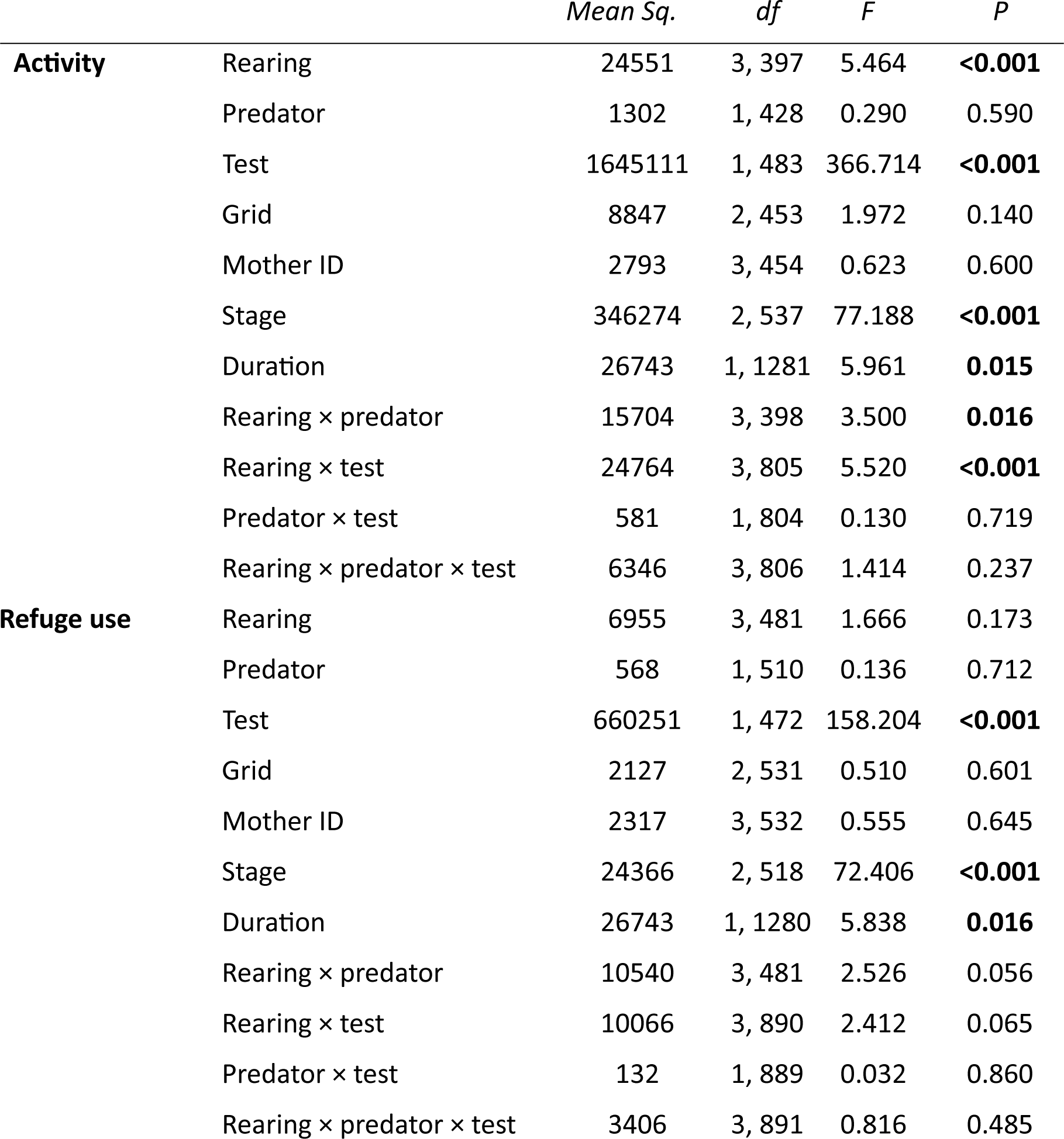
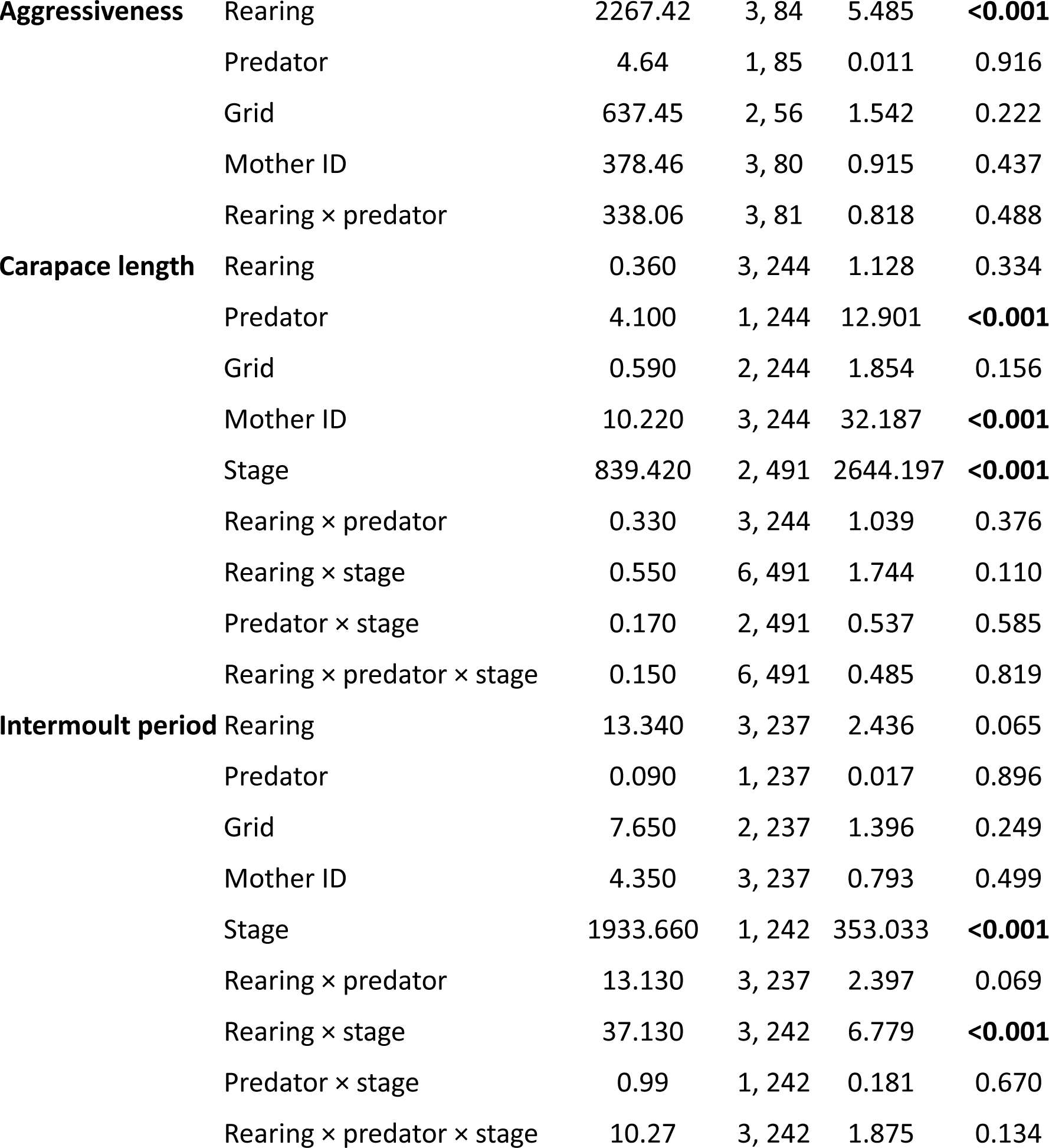
Results from the fixed-factor structure of the generalized linear-mixed models. Rearing treatment (four levels), predator treatment (two levels), test (two levels), grid (four levels), mother ID (four levels), stage (three levels), test duration (in sec), and the interaction between rearing treatment, predator treatment, and test are included as fixed effects in the models for activity and refuge use. In the aggressiveness model rearing treatment, predator treatment, their interaction, grid, and mother ID were the fixed effects. For the life-history traits CL and PI, rearing treatment, predator treatment, stage, their (three way) interaction, grid, mother ID, and stage are included as the fixed effects. Significance was α < 0.05 and significant results are in bold.

|  |  | *Mean Sq.* | *df* | *F* | *P* |
| --- | --- | --- | --- | --- | --- |
| **Activity** | Rearing | 24551 | 3, 397 | 5.464 | **<0.001** |
|  | Predator | 1302 | 1, 428 | 0.290 | 0.590 |
|  | Test | 1645111 | 1, 483 | 366.714 | **<0.001** |
|  | Grid | 8847 | 2, 453 | 1.972 | 0.140 |
|  | Mother ID | 2793 | 3, 454 | 0.623 | 0.600 |
|  | Stage | 346274 | 2, 537 | 77.188 | **<0.001** |
|  | Duration | 26743 | 1, 1281 | 5.961 | **0.015** |
|  | Rearing × predator | 15704 | 3, 398 | 3.500 | **0.016** |
|  | Rearing × test | 24764 | 3, 805 | 5.520 | **<0.001** |
|  | Predator × test | 581 | 1, 804 | 0.130 | 0.719 |
|  | Rearing × predator × test | 6346 | 3, 806 | 1.414 | 0.237 |
| **Refuge use** | Rearing | 6955 | 3, 481 | 1.666 | 0.173 |
|  | Predator | 568 | 1, 510 | 0.136 | 0.712 |
|  | Test | 660251 | 1, 472 | 158.204 | **<0.001** |
|  | Grid | 2127 | 2, 531 | 0.510 | 0.601 |
|  | Mother ID | 2317 | 3, 532 | 0.555 | 0.645 |
|  | Stage | 24366 | 2, 518 | 72.406 | **<0.001** |
|  | Duration | 26743 | 1, 1280 | 5.838 | **0.016** |
|  | Rearing × predator | 10540 | 3, 481 | 2.526 | 0.056 |
|  | Rearing × test | 10066 | 3, 890 | 2.412 | 0.065 |
|  | Predator × test | 132 | 1, 889 | 0.032 | 0.860 |
|  | Rearing × predator × test | 3406 | 3, 891 | 0.816 | 0.485 |
| **Aggressiveness** | Rearing | 2267.42 | 3, 84 | 5.485 | **<0.001** |
|  | Predator | 4.64 | 1, 85 | 0.011 | 0.916 |
|  | Grid | 637.45 | 2, 56 | 1.542 | 0.222 |
|  | Mother ID | 378.46 | 3, 80 | 0.915 | 0.437 |
|  | Rearing × predator | 338.06 | 3, 81 | 0.818 | 0.488 |
| **Carapace length** | Rearing | 0.360 | 3, 244 | 1.128 | 0.334 |
|  | Predator | 4.100 | 1, 244 | 12.901 | **<0.001** |
|  | Grid | 0.590 | 2, 244 | 1.854 | 0.156 |
|  | Mother ID | 10.220 | 3, 244 | 32.187 | **<0.001** |
|  | Stage | 839.420 | 2, 491 | 2644.197 | **<0.001** |
|  | Rearing × predator | 0.330 | 3, 244 | 1.039 | 0.376 |
|  | Rearing × stage | 0.550 | 6, 491 | 1.744 | 0.110 |
|  | Predator × stage | 0.170 | 2, 491 | 0.537 | 0.585 |
|  | Rearing × predator × stage | 0.150 | 6, 491 | 0.485 | 0.819 |
| **Intermoult period** | Rearing | 13.340 | 3, 237 | 2.436 | 0.065 |
|  | Predator | 0.090 | 1, 237 | 0.017 | 0.896 |
|  | Grid | 7.650 | 2, 237 | 1.396 | 0.249 |
|  | Mother ID | 4.350 | 3, 237 | 0.793 | 0.499 |
|  | Stage | 1933.660 | 1, 242 | 353.033 | **<0.001** |
|  | Rearing × predator | 13.130 | 3, 237 | 2.397 | 0.069 |
|  | Rearing × stage | 37.130 | 3, 242 | 6.779 | **<0.001** |
|  | Predator × stage | 0.99 | 1, 242 | 0.181 | 0.670 |
|  | Rearing × predator × stage | 10.27 | 3, 242 | 1.875 | 0.134 |

Chronic exposure to the predator cues had a negative impact on the carapace length of the individuals, with juveniles from the predator treatment being on average smaller than those not exposed to predator cues (Fig. 3a). Overall effects of predator exposure were less evident on the other traits. However, we observed that the interaction between rearing conditions and predator cues explained a large portion of the behavioural variance observed (Figs. 2a, 2b). For instance, juveniles that had access to shelters during development, or that were raised in the presence of both a shelter and substrate, were more active and aggressive than control individuals in the absence of predators (Figs 2a, 2b). On the contrary, effects of the rearing treatment disappeared when individuals were simultaneously exposed to predator cues during their development (Figs 2a, 2b).

**Figure 3.**
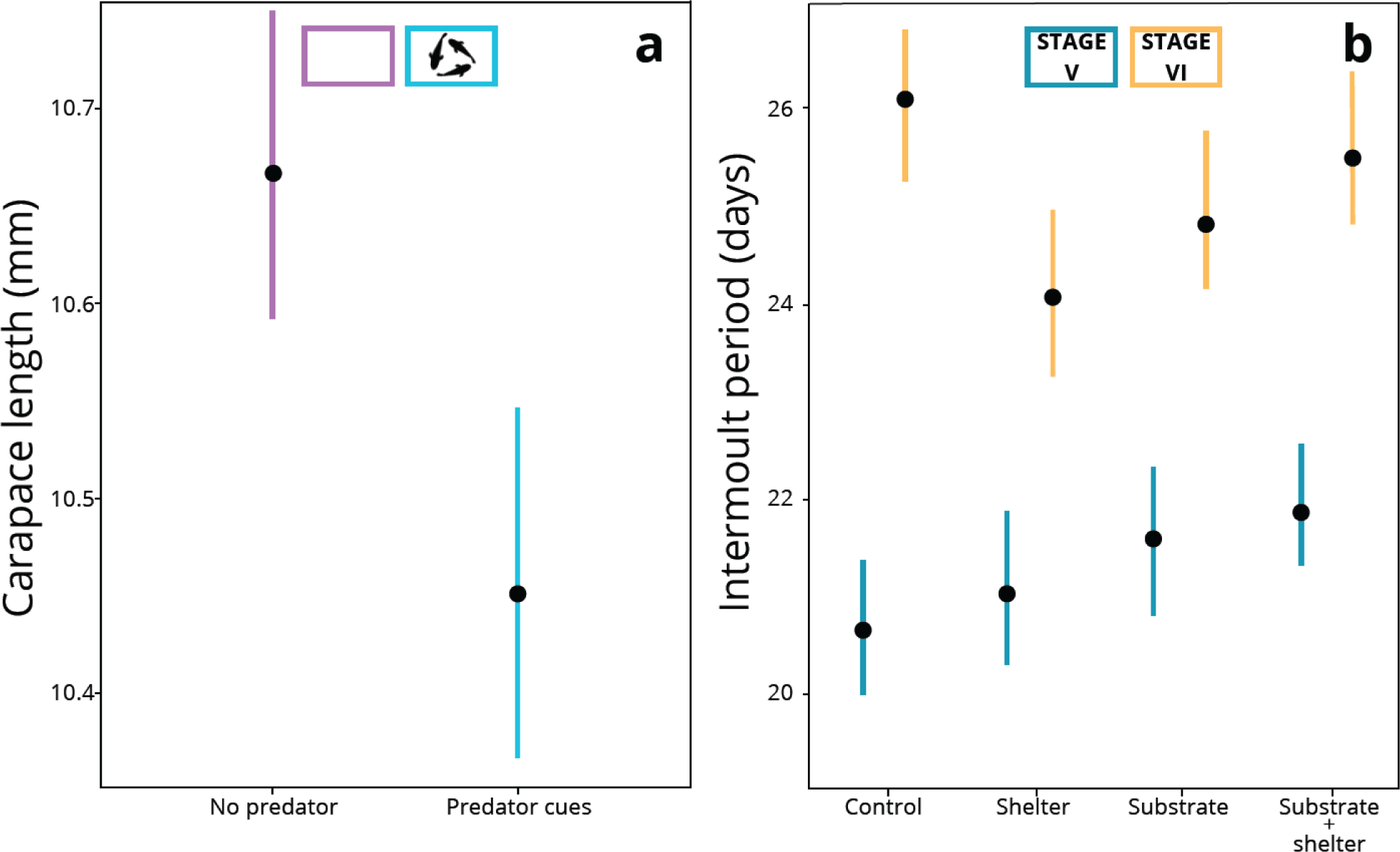
Effects of the different rearing conditions on life-history traits. In (a) is reported the mean effect of predator cues on carapace length, and in (b) the effect of rearing treatment on the intermoult period of the individuals.

The behavioural and life-history traits of the individuals varied, on average, across ontogenetic stages (Table 1; Fig. 3b). For instance, activity levels decreased and refuge use increased over development. We also observed that intermoult period (i.e. growth rate) of the individuals varied between ontogenetic stages depending on the rearing treatment experienced by the individuals (Fig. 3b): the time between two consecutive moults did not differ between rearing treatments early in life, but individuals reared in the presence of substrate and/or a shelter (but not both substrate and shelter together) grew faster than control animals later in life (Fig. 3b). As expected, activity levels and refuge use varied between tests (novel environment, NE, and predator avoidance, PA; Table 1): individuals spent on average less time exploring the experimental arena and hid longer in their refuge when live predators were present (PA) than in the absence of predator cues (NE).

We found evidence that the inclusion of random intercepts (i.e. individual IDs) and test and stage as random slopes increased model fit (Table 2). In other words, individuals differed in behaviour among each other consistently over time (i.e. behavioural individuality), and also differed from each other in the way that they adjusted their activity levels and refuge use across tests and developmental stages (i.e. behavioural plasticity). Instead, no differences were detected between models with rearing treatment, predator treatments, or mother IDs as the random slopes, and we did not retain these terms in our final models (Table 2).

**Table 2.**
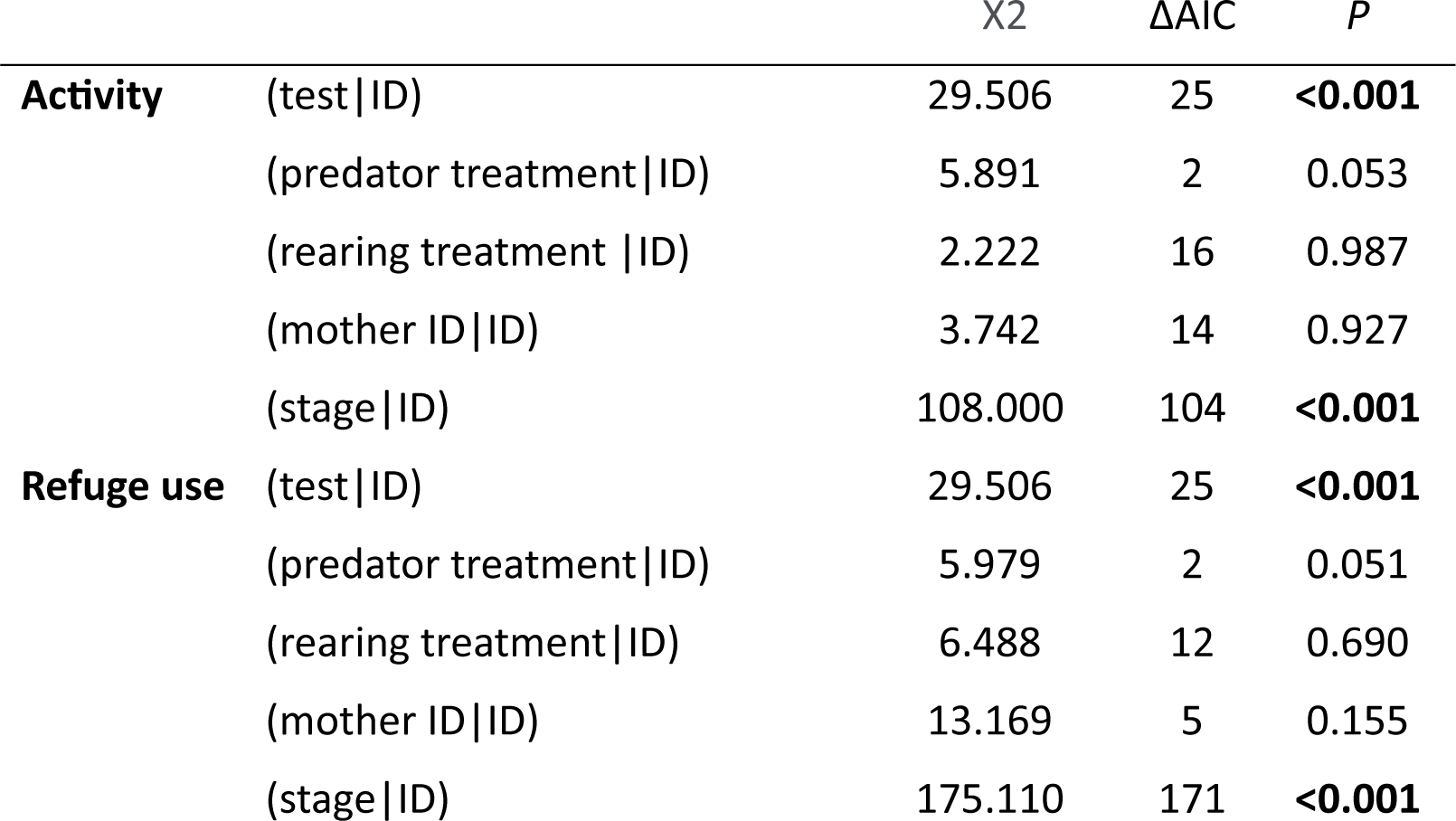
Comparison of LMMs with different random (co)variance structures for activity (time spent in activity) and refuge use (time spent inside the shelter). We compared two models with both random intercepts (ID) and slopes (test, predator treatment, rearing treatment, mother ID, or stage) across individuals against a reduced model with only random intercepts. We also compared the model with random intercepts with a model in which random intercepts were excluded. Significant improvements in the model fit are tested with both Akaike information criteria (ΔAIC) and likelihood-ratio tests (*P*). Rearing treatment (control, substrate, shelter, substrate + shelter), predator treatment (predator cues, no predators), grid (four levels), mother ID (four levels), stage (three levels), test duration (in sec) are included as fixed effects in models considered for activity and refuge use. Significance was set at α < 0.05 and significant results are in bold; the model with random intercepts and test and stage as the random slope was best supported as the most parsimonious model for activity, while random intercepts and stage as the random slopes explained a significant portion of the variance in refuge use. The random structure is written in the syntax of the *lme4 R* package.

## DISCUSSION

Our analysis provides solid evidence that early-life exposure to structural and sensory stimuli in the rearing environment have important effects on both the average and the individual-level behaviours and life-history traits of European lobsters. For instance, the presence of a shelter in the housing structure resulted into higher activity rates of the individuals, while the presence of a gravel substrate reduced their aggressiveness toward intruders. Instead, individuals reared in the presence of either the gravel substrate or the shelter grew faster later in life than control animals, while the exposure to predator cues during development resulted into smaller body sizes of the lobsters.

A main result of this work is that structural enrichments in the rearing enclosures resulted into higher activity levels and lower aggressiveness in the animals. Studies on other aquatic organisms indicate that the presence of environmental enrichments in the rearing enclosures enhances the neural plasticity of these animals by increasing the dopaminergic and serotonergic activity that, in turn, promotes exploratory behaviours, spatial orientation, and learning capacities (Newberry, 1995; Ullah et al., 2017; Arechavala-Lopez et al., 2020). Similarly, structural enrichment was found to reduce aggressiveness in captive-bred fishes (*Tilapia rendaIli*) by Torrezzani and collaborators (2013). In fact, it is reasonable to believe that habitat complexity provides opportunity for individuals to establish territories and safe refugia, thereby reducing basal stress and attenuating their aggressive behaviours. In line with this interpretation, our lobsters raised in the presence of substrate and/or shelter also grew faster (i.e. had lower intermoult periods) later in life, than control groups. Although, to the best of our knowledge, there are no studies that have explored the effects of environmental enrichments on the intermoult period of lobsters, existing research on aquatic organisms indicates that enriched environments reduce stress levels and favour lower resting metabolic rates of the animals, which, in turn, reduce the energy expenditure of the body machinery and favours growth (Howell & Canario, 1987; Millidine et al., 2006; Marcon et al., 2018; Zhang et al., 2021). It is therefore conceivable that environmental enrichment has contributed, over the course of development, to a decrease in stress levels and, consequently, an increase in growth rates, manifested through shorter intermoult periods of the lobsters. Notably, we observed a consistent pattern of longer refuge use and lower activity levels with advancing age, suggesting a shift toward a properly benthic behaviour over development that is well known for these animals (Botero & Atema, 1982; Cobb et al., 1989; Castro & Cobb, 2005; Latini et al., 2023).

Interestingly, our work points out that the exposure to predatory cues during development can result into a smaller body size of the individuals (i.e., carapace length). According with classic life-history theory, in which different predatory regimes result in population having different life-history strategies (Reznick & Endler, 1982), a reduction in body size can be interpreted as a trade-off between securing resources and reducing chances of being predated (Houston et al., 1993; Krause et al., 1998; McNamara et al., 2005). In fact, prior evidence indicates that prey tend to reduce activity levels and increase refuge use in response to the presence of a predator (Lima & Dill, 1990; Sih & McCarthy, 2002). In support of this interpretation, we observed that individuals spent on average more time hiding in the refuge when tested in the presence of their predators (PA test) than when predators were absent (NE test). But effects of the predator treatment were not only related to changes in the life-history traits of the animals, and shaped even the behavioural profiles of our individuals. Specifically, the exposure to predator cues during development reduced the beneficial effects of gravel substrate and shelter on the lobsters’ behaviour: individuals were more active and risk prone (i.e. lower refuge use) when raised with substrate and shelter than conspecifics reared in absence of such structural enrichments, but effects tapered away in animals also exposed to predatory cues during development. These findings suggest that when environmental stimuli are combined, their interaction may lead to effects that do not necessarily correspond to those observed when only one enrichment is present (Näslund, 2021).

Ultimately, our results revealed that individuals consistently displayed distinct behaviour over time (i.e. behavioural individuality), alongside variation in the modulation of their activity levels and refuge use (i.e. behavioural plasticity) depending on the presence/absence of predators in their surrounding environment and developmental age. These findings indicate that the early-life environment plays a crucial role in shaping phenotypic variation within animal groups, and impact their ability to respond to rapid environmental changes (De Gasperin et al., 2019; Zocher et al., 2020). Since phenotypic variation among individuals of a population are likely to increase over development (Polverino et al., 2016), it becomes imperative to carefully plan rearing conditions during juvenile stages of lobsters to foster diversity in the phenotypic traits at adulthood and enhance the likelihood of post-release success. While it remains unknown whether these experimentally-induced phenotypic variations will persist over time and after animals are released in the wild, the literature suggests that individual-level measurements performed in the lab well reflect the natural variation observed among the individuals once released into the wild (Laskowski et al., 2016).

A key goal in conservation aquaculture is producing animals that can effectively survive and reproduce in the wild. We propose that introducing ecological-relevant stimuli in the rearing environment can benefit conservation programs by promoting natural diversity in animal phenotypes, which is essential for thriving under challenging environmental conditions.

## Author contributions

LL, GG, CC, and DC conceived the research project and designed the experiments; LL, GG, and EB performed the experiments; GP and LL analyzed the data; GP and LL discussed and interpreted the results, with contribution from CC and CG; GP and LL wrote the manuscript and all authors provided critical feedback.

## Data Availability

Upon acceptance, both data and the *R* code used for the statistics and figures will be made publicly available via the Figshare repository.

## Acknowledgments

The authors thank Bianca Melita Palmas for helping with the figures’ preparation. The Fondo Europeo per gli Affari Marittimi e la Pesca (FEAMP) supported the restocking actions at the CISMAR. The Department of Ecological and Biological Sciences at the University of Tuscia and the Stazione Zoologica Anton Dohrn co-financed this study. The research project was implemented under the National Recovery and Resilience Plan (NRRP), Mission 4 Component 2 Investment 1.4—call for tender No. 3138 of 16 December 2021, rectified in Decree n.3175 of 18 December 2021 by the Italian Ministry of University and Research, funded by the European Union—Next Generation EU. Project code CN_00000033, Concession Decree No. 1034 of 17 June 2022 adopted by the Italian Ministry of University and Research, CUP J83C22000860007, Project title National Biodiversity Future Centre—NBFC.

